# Sinking rates and export flux of Transparent Exopolymer Particles (TEPs) in an eutrophic coastal sea: a case study in the Changjiang (Yangtze River) Estuary

**DOI:** 10.1101/357053

**Authors:** Shujin Guo, Jun Sun

## Abstract

Transparent exopolymer particles (TEPs) are ubiquitous throughout the oceans, and their sedimentation is considered an efficient biological carbon sink pathway. However, the role TEPs play as a carbon sink in eutrophic coastal seas is not well studied. In order to investigate this issue, two cruises in the spring and summer of 2011 were carried out in the Changjiang (Yangtze River) estuary, a typical eutrophic coastal sea. The concentrations, sinking rates, and export flux of TEPs were studied. TEP concentrations ranged from 40.00 to 1040.00 μg Xeq L^−1^ (mean = 209.70 ± 240.93 μg Xeq L^−1^) in spring and from 56.67 to 1423.33 μg Xeq L^−1^ (mean = 433.33 ± 393.02 μg Xeq L^−1^) in summer. A significant positive correlation between TEP concentrations and chlorophyll (Chl) *a* concentrations was observed. TEP sinking rates ranged from 0.08 to 0.57 m d^−1^ (mean = 0.28 ± 0.14 m d^−1^) in spring and from 0.10 to 1.08 m d^−1^ (mean = 0.34 ± 0.31 m d^−1^) in summer. TEP sinking rates were always higher in the upper layers than in the deeper layers during both seasons. The export flux of TEPs was also calculated, and it ranged from 4.95 to 29.40 mg C m^−2^ d^−1^ in spring (mean = 14.66 ± 8.83 mg C m^−2^ d^−1^) and from 6.80 to 30.45 mg C m^−2^ d^−1^ (mean = 15.71 ± 8.73 mg C m^−2^ d^−1^) in summer. This study is the first study on TEP sinking in the Changjiang (Yangtze River) Estuary, and it confirmed that TEP plays a significant role as a carbon sink in the eutrophic coastal sea.

## 1 Introduction

Transparent exopolymer particles (TEPs) are transparent gel-like particles in the ocean [1] that were first identified and named by Alldredge et al in 1993 [2]. They are formed from polysaccharides that are mainly exuded by phytoplankton cells and bacteria [3–5]. TEPs were largely ignored before 1990s due to the lack of techniques for the visualization and quantification of these particles. They are only visible when stained with the polysaccharide specific dye Alcian Blue [1, 2]. Once a method for their visualization was developed, their high concentrations in the ocean were revealed [6]. It has been reported that TEPs are abundant in marine ecosystems, with concentrations varying between 1 and 8000 ml^−1^ [1].

Although classified as particles, TEPs exhibit gel-like properties, such as a high degree of adhesion, high flexibility, and the ability to swell/shrink depending on environmental conditions [1]. These properties allow TEPs to form aggregates with other particles, such as phytoplankton cells, bacteria, and detritus in the water column [7, 8], potentially enhancing sinking fluxes and stimulating the biological carbon pump in the ocean [9]. The size and abundance of TEPs are on the same order of magnitude as phytoplankton cells. This suggests that they could contribute significantly to the total particulate pool in the ocean [10, 11]. As the C: N ratio of TEP is well above the Redfield ratio [10, 12], sedimentation of TEP is also considered to be an efficient pathway of carbon export in the ocean [10, 13, 14].

Coastal seas receive large amounts of riverine inputs and upwelling of nutrients, sustaining a disproportionate high biological productivity in these areas [15]. Despite their small surface area, coastal seas play an important role in the global oceanic carbon cycle [16, 17]. Overall, they are considered as sinks of atmospheric CO_2_ [15]. In conjunction with the high phytoplankton biomass in the coastal sea, TEP levels are also high in these areas [10, 18–22]. Despite high levels of TEPs in the coastal sea and their important implications for carbon export in these areas, few studies have focused on the sinking dynamics of TEPs and their export flux in the coastal sea. This research gap hampers the comprehensive understanding of the biological pump in coastal seas.

The Changjiang (Yangtze River) estuary, located on the continental shelf at the western rim of the Pacific Ocean, is one of the most eutrophic coastal seas in the world [23]. Annual fluxes of dissolved inorganic silicon, nitrogen, and phosphorus from the Yangtze River into the adjacent coastal sea were 2.22×10^6^, 7.84×10^5^, and 1.51×10^4^ tonnes, respectively [24]. The high nutrient supply results in high phytoplankton biomass in this area, and phytoplankton blooms frequently occur from May to August [25]. Due to the high biological productivity and efficient export of particulate organic matter, the Changjiang (Yangtze River) estuary serves as a net sink of atmospheric CO_2_ [26]. Although several studies have been carried out to study the carbon cycle in the Changjiang (Yangtze River) estuary [26–29], there has been no study that investigates TEP concentrations and their role in the carbon export in this area. For this study, two cruises were carried out in the Changjiang (Yangtze River) estuary in spring and summer, 2011. TEP concentrations were measured, and *in situ* TEP sinking rates were determined. Finally, the carbon export flux of TEPs was calculated. The main goal of this study is to understand the role of TEPs in carbon export in the Changjiang (Yangtze River) estuary. This will hopefully provide useful information for TEP studies in other eutrophic coastal seas around the world.

## 2 Methods

### 2.1 Study area

Two cruises were carried out in the Changjiang (Yangtze River) estuary during the spring and summer of 2011. Five stations on each cruise were selected with various hydrographic conditions to collect the samples and carry out the sinking rates experiments. The sampling stations are shown in Fig 1, and the surface environmental parameters of these stations are presented in Table 1.

**Fig 1.**
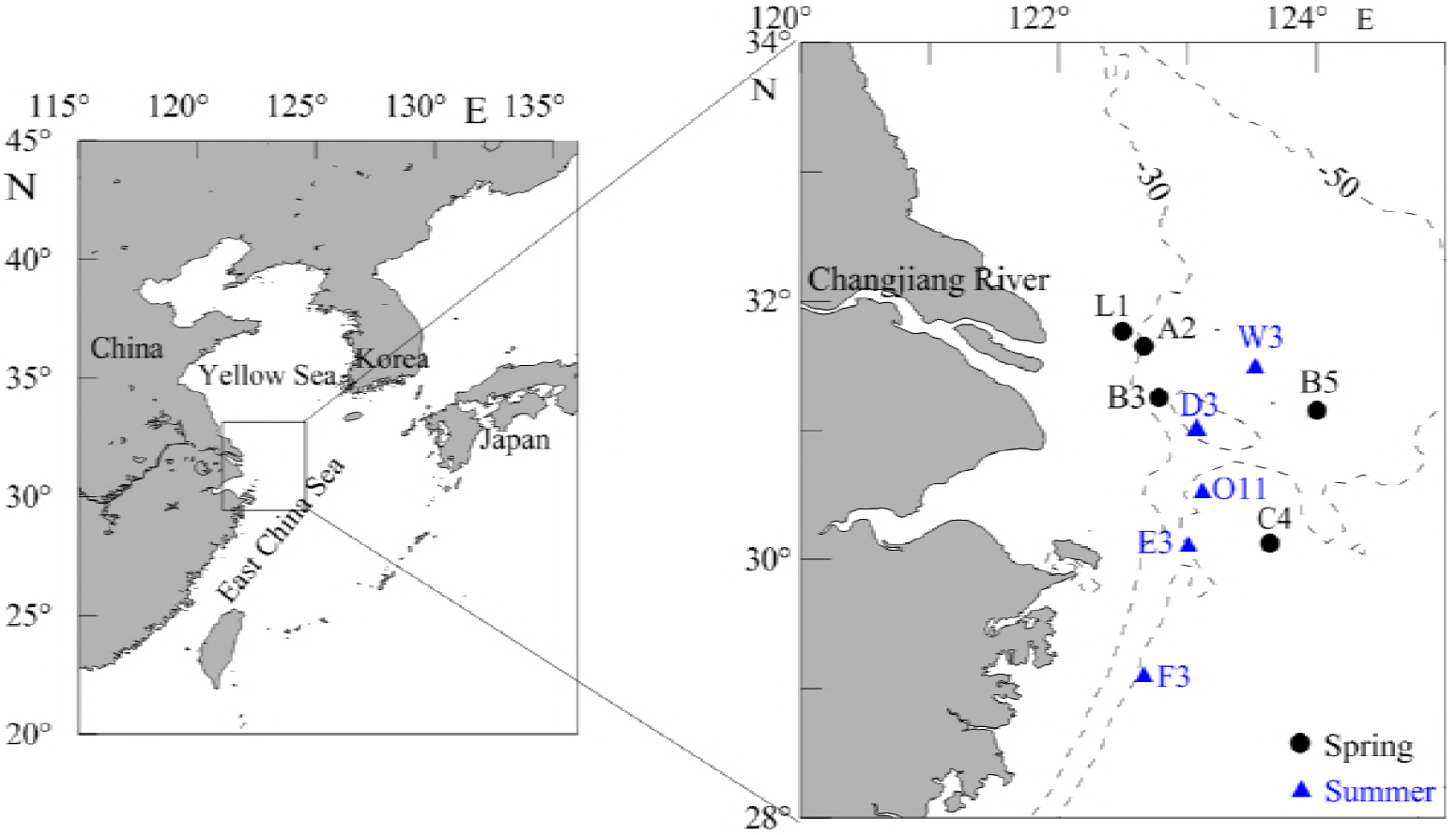
Study sites in the Changjiang (Yangtze River) estuary during spring and summer in 2011.

**Table 1.** The bottom depth (m), temperature (T, °C), salinity (S), chlorophyll *a* (Chl *a*) concentration (μg L^−1^), and concentration (μM) of nutrients (ammonium, nitrate, nitrite, phosphate, and silicate) in the surface layer at the survey stations.

| Cruise | Station | Date | Longitude | Latitude | Bottom depth | T | S | Chl*a* | NH<sub>4</sub><sup>+</sup> | NO<sub>3</sub><sup>-</sup> | NO<sub>2</sub><sup>-</sup> | PO<sub>4</sub><sup>3-</sup> | SiO<sub>3</sub><sup>2-</sup> |
| --- | --- | --- | --- | --- | --- | --- | --- | --- | --- | --- | --- | --- | --- |
| Spring | C4 | 2011-5-22 | 123°39'45" | 30°08'14" | 67.0 | 20.9 | 30.8 | 1.8 | 3.0 | 11.5 | 0.2 | 0.2 | 12.2 |
|  | B5 | 2011-5-25 | 124°00'13" | 31°09'21" | 49.7 | 16.7 | 32.2 | 1.5 | 4.7 | 10.5 | 0.5 | 0.1 | 6.0 |
|  | B3 | 2011-5-25 | 122°46'51" | 31°13'45" | 49.7 | 18.1 | 31.6 | 17.1 | 0.8 | 6.4 | 0.4 | 0.2 | 11.2 |
|  | A2 | 2011-5-25 | 122°38'53" | 31°39'02" | 34.0 | 17.1 | 32.1 | 1.4 | 2.2 | 16.6 | 0.4 | 0.2 | 22.2 |
|  | L1 | 2011-5-31 | 122°28'26" | 31°44'36" | 26.0 | 18.3 | 31.0 | 1.7 | 1.9 | 22.0 | 0.4 | 0.4 | 19.2 |
| Summer | F3 | 2011-8-19 | 122°39'46" | 29°05'53" | 54.0 | 27.7 | 32.0 | 23.8 | 4.6 | 5.2 | 0.7 | 0.9 | 4.7 |
|  | E3 | 2011-8-20 | 123°00'29" | 30°06'12" | 48.4 | 25.2 | 29.5 | 30.2 | 6.9 | 15.0 | 0.8 | 0.2 | 11.3 |
|  | D3 | 2011-8-22 | 123°03'16" | 31°01'15" | 55.3 | 26.2 | 26.0 | 20.0 | 5.6 | 19.1 | 1.0 | 0.3 | 14.9 |
|  | W3 | 2011-8-23 | 123°30'49" | 31°29'18" | 37.8 | 24.8 | 30.4 | 1.5 | 3.2 | 4.5 | 0.5 | 0.2 | 1.4 |
|  | O11 | 2011-8-27 | 123°06'35" | 30°31'13" | 59.4 | 22.0 | 27.5 | 1.8 | 4.4 | 15.5 | 0.6 | 0.4 | 10.3 |

### 2.2 Sampling and analysis

The vertical profiles of temperature (T) and salinity (S) were recorded with a Seabird conductivity, temperature, and depth device (SBE 9/11 plus). Water samples were collected using a rosette sampler at 4 to 5 depths at each station from the surface to the bottom, evenly distributed throughout the water column, for analysis of nutrients, Chl *a*, phytoplankton cell abundance and species composition, and TEP concentration.

#### Nutrient concentrations

Water samples for determination of nutrient concentrations were filtered through acid-cleaned acetate cellulose filters with a 0.45-μm pore size. After being poisoned by HgCl_2_ solution, the filtrates were stored at 0-4°C in the dark until analysis. In the laboratory, nutrients (NO_3_^-^, NO_2_^-^, PO_4_^3-^, SiO_3_^2-^, and NH_4_+) were evaluated with an autoanalyzer (model: SkalarSAN^plus^, Skalar Analysis, Netherlands) according to the manual methods [30].

#### Chl a

Samples for Chl *a* concentration determination were filtered onto 25 mm GF/F filters (Whatman™) and then stored at −20°C in the dark until analysis. Chl *a* was extracted with 90% acetone for 24 h at −20°C in the dark, and samples were then analyzed with a Turner-Designs Trilogy™ laboratory fluorometer [31].

#### Phytoplankton cell abundance and species composition

The samples for phytoplankton analysis were preserved with 2% buffered formalin on board the vessel. In the laboratory, phytoplankton cells were identified and enumerated with an inverted microscope (Olympus, Japan) at a magnification of 200× or 400× according to the Utermöhl method [32].

#### TEP concentrations

Concentrations of TEPs were measured with the methods of Passow and Alldredge [6]. Six 50ml or 100 ml subsamples were vacuum filtered (<0.2 bar) with polycarbonate filters (Millipore; 25 mm diameter; 0.20 μm). Filters were first stained for <5 s with 1 mL of 0.02% Alcian Blue 8GX in 0.06% acetic acid (pH 2.5), and then samples were rinsed with 3 mL of deionized water. Alcian Blue-stained material in the filters was extracted with 6 mL of 80% sulfuric acid for 2 h on an oscillator, and the absorbance of the extracted material was then measured spectrophotometrically at 787 nm. TEPs were quantified by a standard curve prepared with xanthan gum (XG) particles as described by Passow and Alldredge in 1995 [6], and concentrations are expressed in micrograms of XG equivalents per liter (μg Xeq L^−1^).

#### Sinking rates of TEPs

Sinking rates of TEPs were determined at each station, and the SETCOL method [33] was used to measure the sinking rates. For analysis, a Plexiglass column (height = 0.45 m and volume = 750 ml) was filled completely with a homogeneous water sample within 10 min after sampling, and a cover was then placed on the set-up. The Plexiglass column was allowed to settle undisturbed for 2-3 hours aboard the vessel, and the temperature was maintained by pumping water from a thermostatically controlled water bath with water jackets. The settlement experiment was terminated by successively draining the upper, middle, and bottom compartments of the Plexiglass column via taps in the wall of column. The TEP biomass was measured before and after the settlement in all three compartments. These measurements were combined to calculate the sinking rate of TEPs according to the formula:

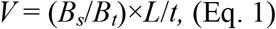

where *V* = sinking rate; *B*_*s*_ = the biomass of TEP settled into the bottom compartment; *B*_*t*_ = the total biomass of TEPs in the column; *L* = length of the column; and *t* = settling interval. Three replicates of the settlement columns were filled with seawater collected from each sampling depth, and the mean of the three sinking rate values was calculated to represent the sinking rate at a particular sampling depth.

### 2.3 Data analysis and calculation

The dominance of phytoplankton species was described by the dominance index (*Y*):

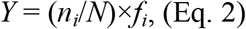

where *n*_*i*_ is the sum of cell abundance for species *i* in all samples; *N* is the sum of cell abundance for all species; and *f*_*i*_ is the frequency of occurrence for species *i* in all samples [34]. SPSS 14.0 was applied to carry out the Pearson Correlation Analysis between TEP concentrations and various environmental parameters.

Carbon flux of TEP estimates were provided by the product of the SETCOL-determined sinking rates mentioned above and the TEP carbon concentrations at the bottom layer. TEP-carbon (*C*_*TEP*_, μg C L^−1^) was calculated with the slope (0.75) from the equation as follows [10]:

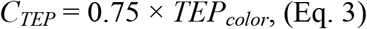

where *TEP*_*color*_ is the TEP concentration with the unit of μg Xeq L^−1^.

## 3 Results

### 3.1 Hydrographic conditions

Vertical profiles of temperature, salinity, and Chl *a* concentration at each station are presented in Fig 2. Surface temperatures ranged from 16.7 to 20.9 °C (mean = 18.2 ± 1.5 °C) in spring and from 22.0 to 27.7 °C (mean = 25.2 ± 1.9 °C) in summer. Surface salinity ranged from 30.8 to 32.2 (mean = 31.5 ± 0.6) in spring and from 26.0 to 32.0 (mean = 29.1 ± 2.1) in summer. Generally, the upper layers were dominated by warm and low salinity water and the deeper layers were dominated by cool and high salinity water during both seasons. High Chl *a* concentrations were always observed in the upper layers at each station, and Chl *a* concentrations exceeded 15 μg L^−1^ in the surface layer at station B3 in spring as well as stations F3, E3, and D3 in summer. At other stations, Chl *a* concentrations were relatively low. Dominant phytoplankton species varied greatly among the two cruises (Table 2). In spring, *Prorocentrum dentatum* was the most dominant species, and cell abundance exceeded 10^6^ cells L^−1^ in the surface layer at station B3. In summer, *Skeletonema* cf. *costatam* was the most dominant species, and cell abundance in the surface layer at stations F3, E3, and D3 exceeded 10^6^ cells L^−1^. Generally, the phytoplankton community in the study area was dominated by dinoflagellates in spring and diatoms in summer.

**Fig 2.**
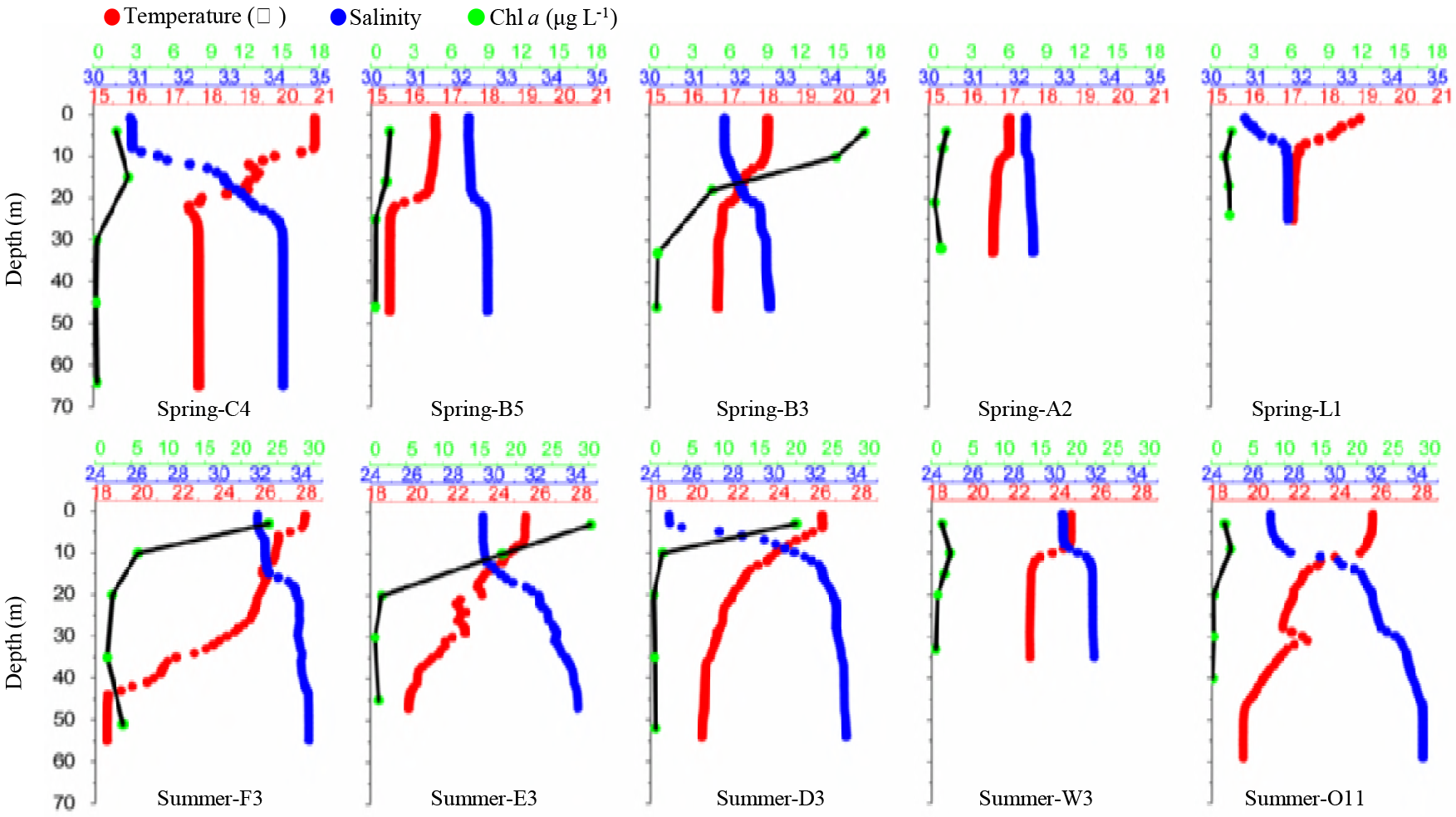
Vertical profiles of temperature, salinity, and Chl *a* concentrations at each station during the two cruises.

**Table 2.**
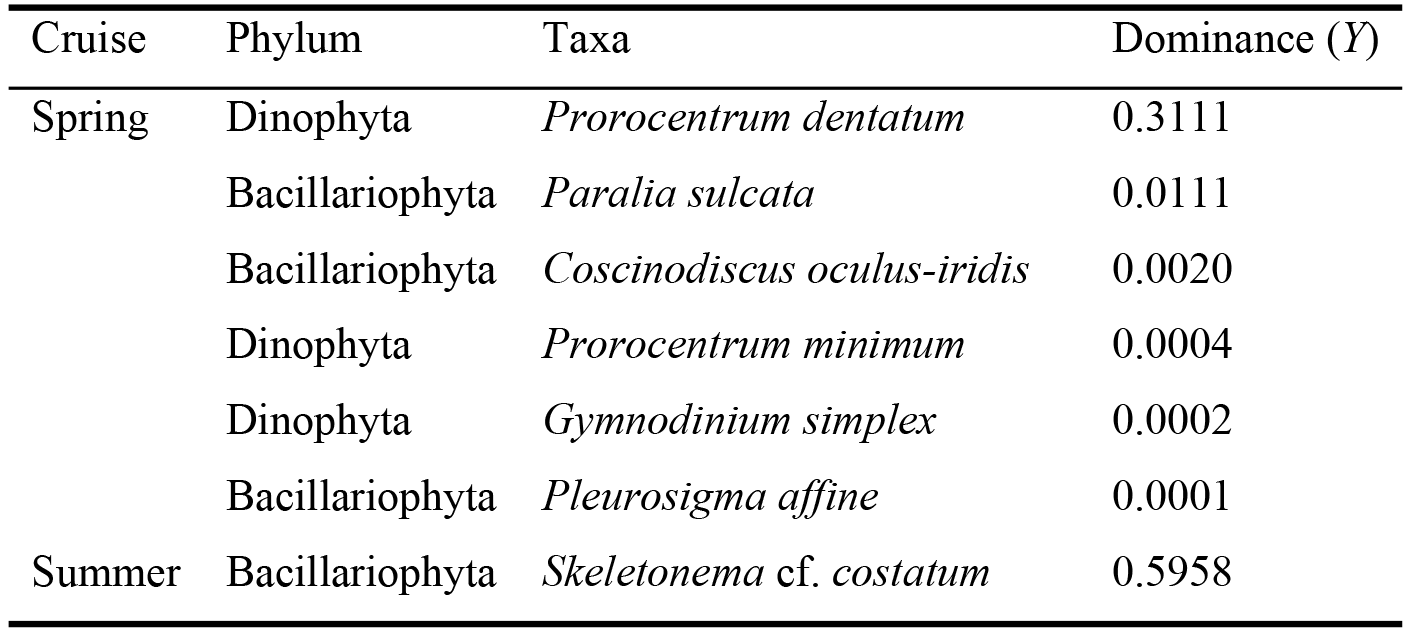
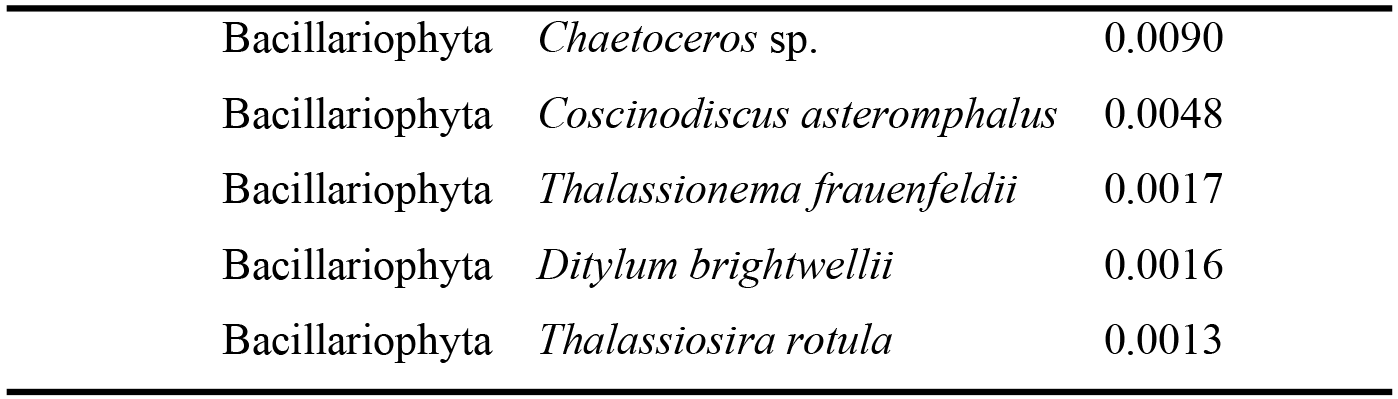
Dominant phytoplankton species in the study area during the two cruises.

| Cruise | Phylum | Taxa | Dominance (*Y*) |
| --- | --- | --- | --- |
| Spring | Dinophyta | *Prorocentrum dentatum* | 0.3111 |
|  | Bacillariophyta | *Paralia sulcata* | 0.0111 |
|  | Bacillariophyta | *Coscinodiscus oculus-iridis* | 0.0020 |
|  | Dinophyta | *Prorocentrum minimum* | 0.0004 |
|  | Dinophyta | *Gymnodinium simplex* | 0.0002 |
|  | Bacillariophyta | *Pleurosigma affine* | 0.0001 |
| Summer | Bacillariophyta | *Skeletonema cf. costatum* | 0.5958 |

|  |  |  |
| --- | --- | --- |
| Bacillariophyta | *Chaetoceros* sp. | 0.0090 |
| Bacillariophyta | *Coscinodiscus asteromphalus* | 0.0048 |
| Bacillariophyta | *Thalassionema frauenfeldii* | 0.0017 |
| Bacillariophyta | *Ditylum brightwellii* | 0.0016 |
| Bacillariophyta | *Thalassiosira rotula* | 0.0013 |

### 3.2 TEP concentrations

Concentrations of TEPs at each station are shown in Fig 3. TEP concentrations ranged from 40.00 to 1040.00 μg Xeq L^−1^ with an average value of 209.70 μg Xeq L^−1^ in spring, and they ranged from 56.67 to 1423.33 μg Xeq L^−1^ in summer with an average value of 433.33 μg Xeq L^−1^. In the surface layer, TEP concentrations ranged from 173.33 to 840.00 μg Xeq L^−1^ in spring, and from 473.33 to 1423.33 μg Xeq L^−1^ in summer. High values appeared at station B3 in spring as well as stations F3, E3, and D3 in summer. High TEP concentrations always appeared in upper layers at each station, and they decreased as water depth increased during both cruises. TEP concentration showed significant positive correlations with Chl *a* concentration and temperature in the study area (Table 3), and no other environmental parameter was found to have a significant correlation with TEP concentration.

**Fig 3.**
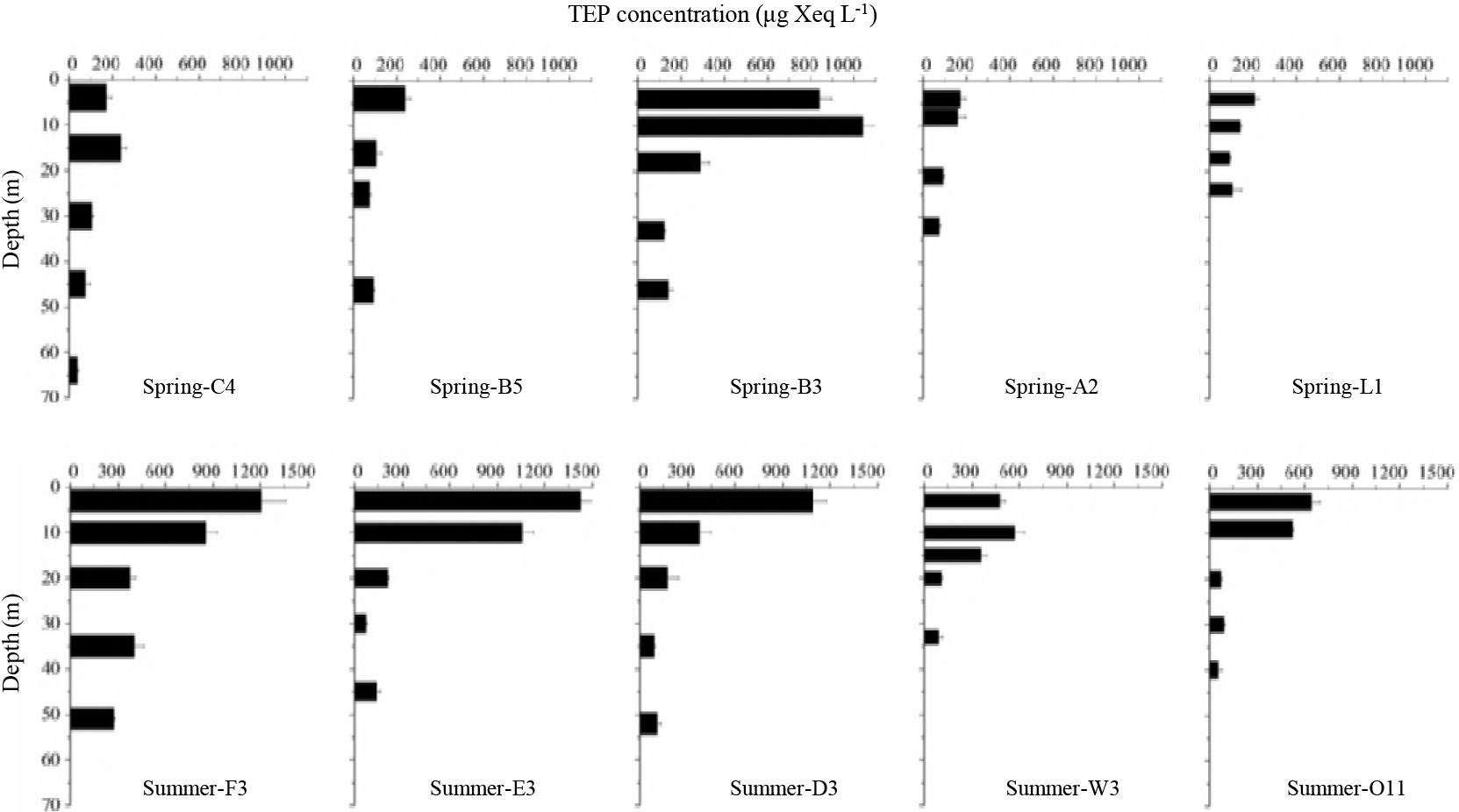
TEP concentrations (μg Xeq L^−1^) at the survey stations during the two cruises.

**Table 3.** Correlation analysis between TEP concentrations and environmental parameters at the survey stations.

| Cruise |  | Temperature | Salinity | Chl *a* | NH<sub>4</sub><sup>+</sup> | NO<sub>2</sub><sup>-</sup> | NO<sub>3</sub><sup>-</sup> | PO<sub>4</sub><sup>3-</sup> | SiO<sub>3</sub><sup>2-</sup> |
| --- | --- | --- | --- | --- | --- | --- | --- | --- | --- |
| Spring | *R* | -.091 | .111 | .794** | -.586 | .209 | -.660 | .053 | -.316 |
| Summer | *R* | .491 | .092 | .870** | .872 | .721 | .304 | .240 | .473 |
| All | *R* | .726* | -.397 | .849** | .747 | .803 | -.151 | .435 | -.274 |

\* indicates $P < 0.05$ ; \*\* indicates $P < 0.01$ .

### 3.3 Sinking rates of TEPs

Sinking rates of TEPs are shown in Fig 4. Sinking rates of TEPs ranged from 0.08 to 0.57 m d^−1^ in spring with an average value of 0.28 m d^−1^ and from 0.10 to 1.08 m d^−1^ in summer with an average value of 0.34 m d^−1^. Except for the surface layer at station F3 in summer, most of the sinking rates determined in the study area were below 1 m d^−1^. For the surface layer in spring, the sinking rate of TEPs at station B3 was the highest, followed by stations B5, L1, and A2. Station C4 exhibited the lowest TEP sinking rate for the surface layer in spring. For the surface layer in summer, sinking rates of TEPs were obviously higher at stations F3, E3, and D3 than at stations W3 and O11. Vertically, sinking rates of TEPs were higher in the upper layers than in the deeper layers during both cruises (Fig 5).

**Fig 4.**
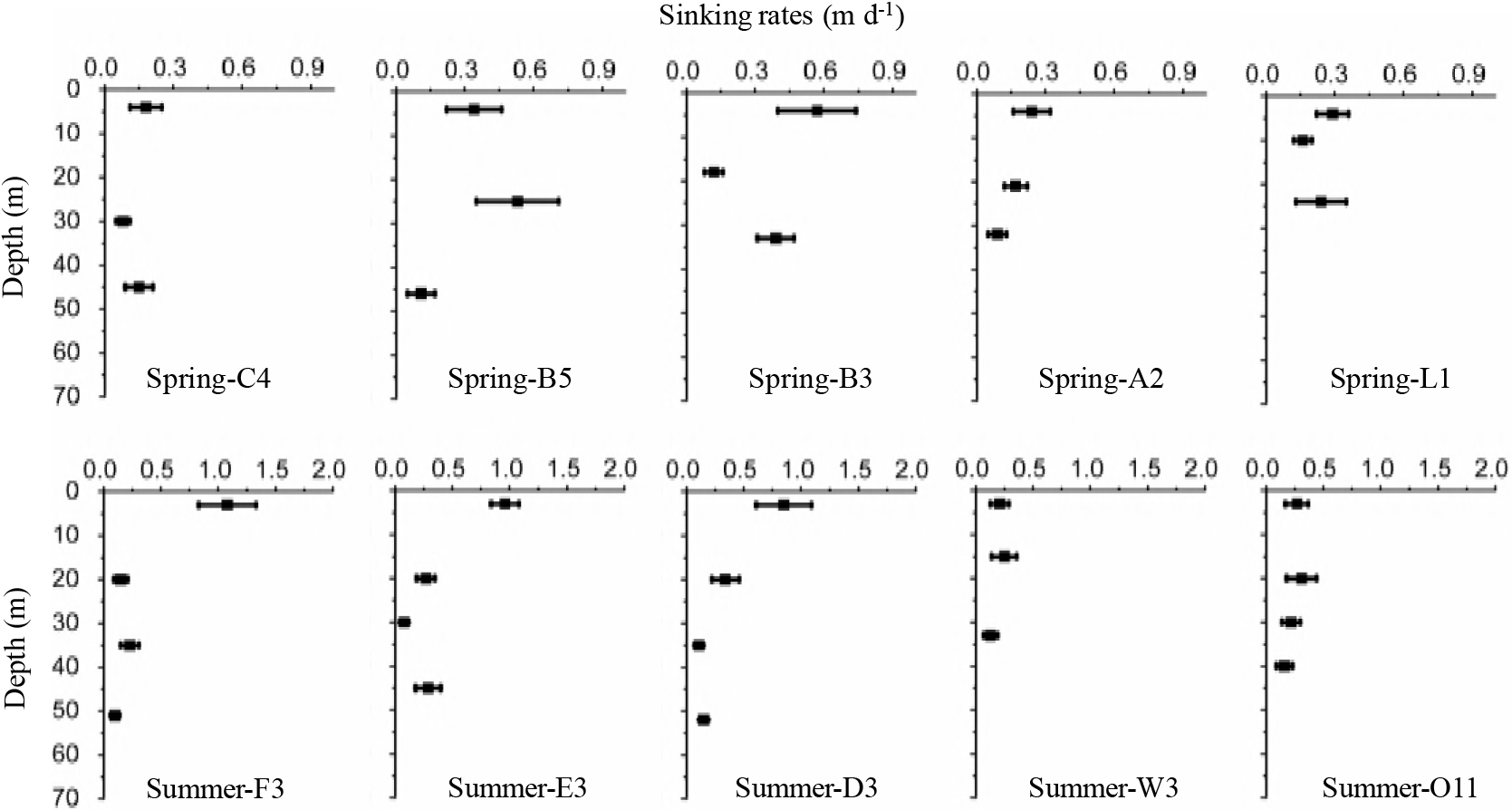
Sinking rates of TEPs at each station during the two cruises as measured with the SETCOL method.

**Fig 5.**
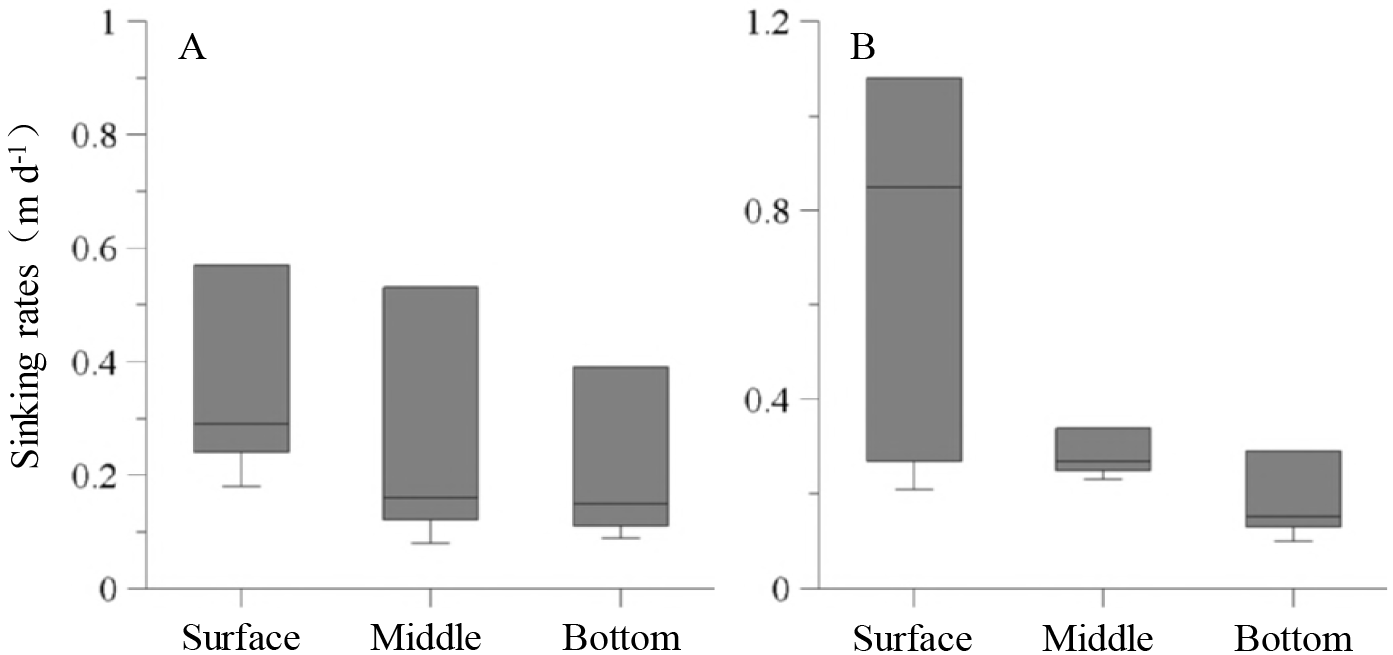
Comparison of TEP sinking rates at the surface, middle, and bottom layer during (A) spring and (B) summer in 2011.

### 3.4 Export flux of TEPs

The estimated export flux of TEPs at each station is shown in Fig 6. In spring, export flux of TEPs ranged from 4.95 to 29.40 mg C m^−2^ d^−1^ with an average value of 14.66 ± 8.83 mg C m^−2^ d^−1^, and the export flux at station B3 was obviously higher than that at the other four stations. In summer, export flux of TEPs ranged from 6.80 to 30.45 mg C m^−2^ d^−1^ with an average value of 15.71 ± 8.73 mg C m^−2^ d^−1^, and the export flux at stations F3 and E3 were obviously higher than that at the other three stations. Generally, the TEP export flux was similar between spring and summer.

**Fig 6.**
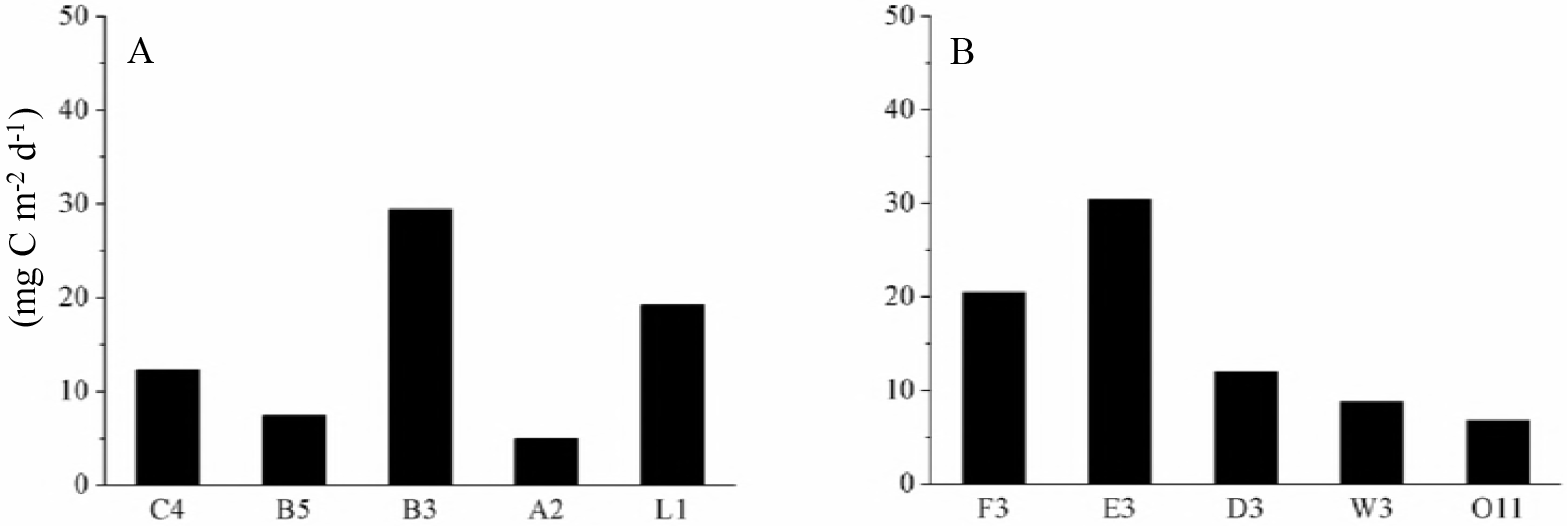
Export flux of TEPs at each station in (A) spring and (B) summer as estimated by SETCOL-measured sinking rates.

## 4 Discussion

### 4.1 TEP concentration in the Changjiang (Yangtze River) estuary and its relationship with environmental parameters

TEP concentrations ranged from 40.00 to 1423.33 μg Xeq L^−1^ in the study area, which is within the range reported in other coastal seas around the world (Table 4). TEP concentrations in the 10 m layer at station B3 during spring and within the surface layer at stations F3, E3, and D3 during summer were high, exceeding 1000 μg Xeq L^−1^. Phytoplankton biomasses at these stations were also high. A *P. dentatum* bloom was observed at station B3 in spring, and a *S.* cf. *costatum* bloom was observed at stations F3, E3 and D3 in summer. Chl *a* concentrations in the upper layers at these stations were also obviously higher than those at the other stations (Fig 2). Several studies have observed high TEP concentrations during phytoplankton blooms in coastal seas around the world [8, 35–39]. The coincident maxima for TEPs and phytoplankton biomass in these areas are consistent with the concept that TEPs are mainly produced by growing and senescing phytoplankton cells [1, 13]. The significant positive correlation between TEP concentrations and Chl *a* concentrations in this study provides further support for this concept (Table 3).

**Table 4.**
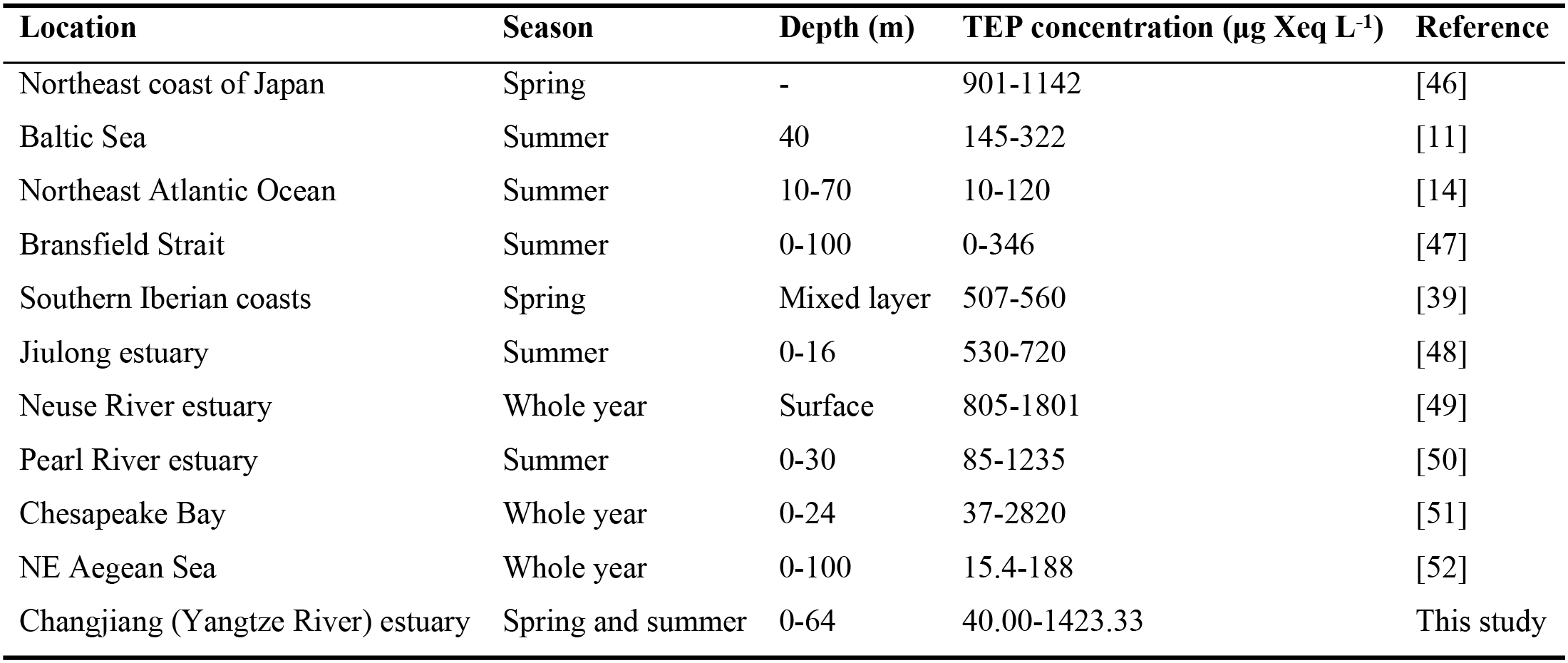
TEP concentrations in the Changjiang (Yangtze River) estuary and other coastal seas around the world.

| Location | Season | Depth (m) | TEP concentration ($\mu\text{g Xeq L}^{-1}$) | Reference |
| --- | --- | --- | --- | --- |
| Northeast coast of Japan | Spring | - | 901-1142 | [46] |
| Baltic Sea | Summer | 40 | 145-322 | [11] |
| Northeast Atlantic Ocean | Summer | 10-70 | 10-120 | [14] |
| Bransfield Strait | Summer | 0-100 | 0-346 | [47] |
| Southern Iberian coasts | Spring | Mixed layer | 507-560 | [39] |
| Jiulong estuary | Summer | 0-16 | 530-720 | [48] |
| Neuse River estuary | Whole year | Surface | 805-1801 | [49] |
| Pearl River estuary | Summer | 0-30 | 85-1235 | [50] |
| Chesapeake Bay | Whole year | 0-24 | 37-2820 | [51] |
| NE Aegean Sea | Whole year | 0-100 | 15.4-188 | [52] |
| Changjiang (Yangtze River) estuary | Spring and summer | 0-64 | 40.00-1423.33 | This study |

TEP formation and distribution are controlled by several physical and biological factors, including temperature [40, 41], salinity [42], nutrient levels [43, 44], and phytoplankton species composition [1]. In this study, TEP concentrations showed a significant positive correlation with temperature (Table 3). Claquin et al. (2008) studied the effects of temperature on photosynthetic parameters and TEP production in eight species of marine microalgae, and they found that temperature influenced TEP production by affecting the photosynthetic activity of phytoplankton cells [40]. Fukao et al. (2012) studied the effects of temperature on cell growth and production of TEPs by the diatom *Coscinodiscus granii*, and they found higher growth rates of *C. granii* at higher temperatures [41]. This is likely responsible for the high production of TEPs at higher temperature. Therefore, temperature impacts TEP production by affecting phytoplankton physiological activity. In this study, salinity and nutrient concentrations showed no significant correlations with TEP concentration (Table 3), despite their reported effect on TEP formation [42–45]. The limited sample data and insignificant variation in salinity and nutrient concentrations among sampling stations (Table 1) may be responsible for the poor correlations between TEPs and these environmental parameters in this study.

### 4.2 TEP sinking rates in the Changjiang (Yangtze River) estuary and comparisons with other studies

The TEP sinking rates as determined by the SETCOL method represent the settling of TEPs in the water column in the absence of turbulence. The real sinking rates of TEPs should be the sum of the intrinsic sinking rates of TEPs and the instantaneous rate of motion in seawater. The effect of turbulence on the sinking rates of micro-sized particles in the ocean was shown to be complex [53, 54], and the mechanisms underlying how turbulence affects the sinking rates of these particles remains unclear. Regardless of the level of turbulence, the SETCOL-measured sinking rate is an important parameter for understanding the movement of TEPs in seawater. Due to the simple technology and reliable determination of results, the SETCOL method has also been used to measure sinking rates of TEP in other studies [55, 56].

The 0.08-1.08 m d^−1^ range of TEP sinking rates measured with the SETCOL method in this study fell within the range reported in other studies (Table 5). It has been reported that TEPs are less dense than seawater, with an estimated density between 0.70 to 0.84 g cm^−3^ [55]. Therefore, ballast-free, ‘pure’ TEPs would ascend in the seawater. This is consistent with the results of Azetsu-Scott and Passow (2004) [55] and Mari (2008) [56] that the sinking rates of TEPs could be negative (Table 5). However, ballast-free TEPs are unlikely to exist in large numbers in coastal seas due to high concentrations of suspended inorganic and organic particulate matter in these areas. As TEPs are extremely sticky [37, 58], they can form aggregates with ambient phytoplankton cells, bacteria, mineral clays, and detritus [38]. This aggregation probably increases the weight of TEPs and allows them to sink to deeper waters [59]. Vicente et al. (2009) calculated TEP sinking rates in an oligotrophic reservoir via sediment trap results [57]. They found that TEP sinking rates ranged from 1.12 to 1.31 m d^−1^, which is slightly higher than our results. In the Vincente study, they found phytoplankton aggregates in the sediment traps, and these aggregates containing TEPs are likely responsible for the higher TEP sinking rates reported in their study. However, these large phytoplankton aggregates containing TEPs might be lost during the discrete sampling of the SETCOL method. Therefore, this study underestimated TEP sinking rates by excluding the effect of large phytoplankton aggregates.

**Table 5.** Comparison of the TEP sinking rates in this study with results from other studies.

| Location | TEP form | Sinking rate (m d<sup>-1</sup>) | Reference |
| --- | --- | --- | --- |
| Seawater from Santa Barbara Channel | Particle-free TEPs | -0.22-0.04 | [55] |
| Water from New Caledonia | Aggregates of TEPs and latex beads | -0.29-0.49 | [56] |
| Oligotrophic reservoir (Quéntar) | Natural TEPs | 1.12-1.31 | [57] |
| Changjiang (Yangtze River) estuary | Natural TEPs | 0.08-1.08 | This study |

### 4.3 Potential role of TEPs in carbon export within the Changjiang (Yangtze River) estuary

It has been reported that TEPs disappear from the euphotic zone via two main pathways: degradation by bacteria and sinking processes associated with other particles [39]. Researchers have concluded that the former pathway is less important than the sinking process due to the refractory nature of TEPs [3, 60]. Therefore, sedimentation of TEPs represents the dominant pathway of their removal in the ocean. As the concentration and carbon content of TEPs is sometimes within the same order of magnitude as that of phytoplankton cells [1, 13], export of carbon via sedimentation of TEPs is significant. In the Santa Barbara Channel, the sedimentation flux of TEPs at 500 m was found to range from 7 to 70 mg C m^−2^ d^−1^, contributing roughly 30% to the particulate organic carbon (POC) flux in this area [13]. In an oligotrophic reservoir in southern Spain, the sedimentation flux of TEPs ranged from 0.51 to 177.04 mg C m^−2^ d^−1^ at the bottom layer, contributing between 0.02% and 31% to the carbon export to sediments [59]. In this study, the export flux of TEPs was calculated within the range of 4.95 to 29.40 mg C m^−2^ d^−1^ in spring and 6.80 to 30.45 mg C m^−2^ d^−1^ in summer (Fig 6). It is unfortunate that no data regarding the export flux of POC was obtained for this study. However, Guo et al studied the export flux of phytoplankton cells in the Changjiang (Yangtze River) estuary during the spring and summer of 2011, and they found that the export flux of phytoplankton cells was 26.10 ± 26.25 mg C m^−2^ d^−1^ in spring and 63.13 ± 48.16 mg C m^−2^ d^−1^ in summer [29]. These levels are slightly higher than the levels of TEPs export flux in this study. Because phytoplankton cell sedimentation is always considered an important pathway of carbon export in the coastal sea [61, 62], the similarity between the TEP export in this study and the phytoplankton cell export indicates that sedimentation of TEPs should also be an important pathway of POC export in the Changjiang (Yangtze) River estuary. As TEPs are types of transparent gel-like particles found in seawater, they have been largely ignored and have received much less attention in studies on POC export when compared with phytoplankton cells and zooplankton fecal pellets [61, 62]. The results of this study suggest that they should be taken into account when studying sinking POC in the chanjiang (Yangtze River) estuary.

## 5 Conclusion

TEP concentrations, sinking rates, and export flux were studied in the Changjiang (Yangtze River) estuary during the spring and summer of 2011. TEP concentrations ranged from 40.00 to 1040.00 μg Xeq L^−1^ in spring and from 56.67 to 1423.33 μg Xeq L^−1^ in summer. TEP concentrations exhibited a significant positive correlation with Chl *a* concentrations. TEP sinking rates determined via the SETCOL method ranged from 0.08 to 0.57 m d^−1^ in spring and from 0.10 to 1.08 m d^−1^ in summer, and these rates were always higher in the upper layers than those in the deeper layers. The export flux of TEPs was also calculated, and it was slightly lower than that of phytoplankton cells in the study area. This study is the first of its kind conducted in the Changjiang (Yangtze River) estuary, and it confirmed that TEPs play a significant role in the carbon export in this area. Due to their transparency, TEPs have been largely ignored in previous studies of POC export in the Changjiang (Yangtze River) estuary. Future studies should pay additional attention to this important component of POC.

## Acknowledgements

We thank the crew and captain of the R/V *Shiyan3* and R/V *Beidou* for the logistic support during the cruise. We also thank professor Sumei Liu for providing the nutrients data and professor Daji Huang for providing the temperature and salinity data. This study was supported by the National Basic Research Program of China (No. 2015CB954002 and 2011CB409804), the National Natural Science Foundation of China (41676112, 4127612, 491751202 and 31700425), the Science Fund for University Creative Research Groups in Tianjin (TD12-5003) and the Changjiang Scholar Program of Chinese Ministry of Education of China to Jun Sun.

## References

1. Passow U. Transparent exopolymer particles (TEP) in aquatic environments. Progress in Oceanography. 2002; 55: 287–333. https://doi.org/10.1016/S0079-6611(02)00138-6

2. Alldredge AL, Passow U, Logan BE. The abundance and significance of a class of large, transparent organic particles in the ocean. Deep-Sea Research I. 1993; 40: 1131–1140. https://doi.org/10.1016/0967-0637(93)90129-Q

3. Stoderegger K, Herndl GJ. Production of exopolymer particles by marine bacterioplankton under contrasting turbulence conditions. Marine Ecology Progress Series. 1999; 189: 9–16. https://doi.org/10.3354/meps189009

4. García CM, Prieto L, Vargas M, Echevarría F, García-Lafuente J, Ruiz J. Hydrodynamics and the spatial distribution of plankton and TEP in the Gulf of Cádiz (SW Iberian Peninsula). Journal of Plankton Research.2002; 24: 817–833. https://doi.org/10.1093/plankt/24.8817

5. Passow U. Production of transparent exopolymer particles (TEP) by phyto- and bacterioplankton. Marine Ecology Progress Series. 2002; 236: 1–12. https://doi.org/10.3354/meps236001

6. Passow U, Alldredge AL. A dye-binding assay for the spectrophotometric measurement of transparent exopolymer particles (TEP). Limnology and Oceanography. 1995; 40(7): 1326–1335. https://doi.org/10.4319/lo.1995.40J.1326

7. Simon M, Grossart HP, Schweitzer B, Ploug H. Microbial ecology of organic aggregates in aquatic ecosystems. Aquatic Microbial Ecology. 2002; 28: 175–211. https://doi.org/10.3354/ame028175

8. Bar-Zeev E, Berman T, Rahav E, Dishon G, Herat B, Berman-Frank I. Transparent exopolymer particle (TEP) dynamics in the eastern Mediterranean Sea. Marine Ecology Progress Series. 2011; 431: 107–118. https://doi.org/10.3354/meps09110

9. Koeve W. Magnitude of excess carbon sequestration into the deep ocean and the possible role of TEP. Marine Ecology Progress Series. 2005; 291: 53–64. https://doi.org/10.3354/meps291053

10. Engel A, Passow U. Carbon and nitrogen content of transparent exopolymer particles (TEP) in relation to their Alcian Blue adsorption. Marine Ecology Progress Series. 2001; 219: 1–10. https://doi.org/10.3354/meps219001

11. Engel A. Direct relationship between CO2 uptake and transparent exopolymer particles production in natural phytoplankton. Journal of Plankton Research. 2002; 24(1): 49–53. https://doi.org/10.1093/plankt/24.L49

12. Mari X. Carbon content and C: N ratio of transparent exopolymeric particles (TEP) produced by bubbling exudates of diatoms. Marine Ecology Progress Series. 1999; 183: 59–71. https://doi.org/10.3354/meps183059

13. Passow U, Shipe RF, Murray A, Pak DK, Brzezinski MA, Alldredge AL. The origin of transparent exopolymer particles (TEP) and their role in the sedimentation of particulate matter. Continental Shelf Research. 2001; 21(4): 327–346. https://doi.org/10.1016/S0278-4343(00)00101-1

14. Engel A. Distribution of transparent exopolymer particles (TEP) in the northeast Atlantic Ocean and their potential significance for aggregation processes. Deep Sea Research I. 2004; 51(1): 83–92. https://doi.org/10.1016/j.dsr.2003.09.001

15. Chen CTA, Borges AV. Reconciling opposing views on carbon cycling in the coastal ocean: continental shelves as sinks and nearshore ecosystems as sources of atmospheric CO2. Deep-Sea Research II. 2009; 56: 578–590. https://doi.org/10.1016/j.dsr2.2009.01.001

16. Cai WJ. Estuarine and coastal ocean carbon paradox: CO_2_ sinks or sites of terrestrial carbon incineration? Annual Review of Marine Science. 2011; 3: 123–145. https://doi.org/10.1146/annurev-marine-120709-142723

17. Bauer JE, Cai WJ, Raymond PA, Bianchi TS, Hopkinson CS, Regnier PAG. The changing carbon cycle of the coastal ocean. Nature. 2013; 504: 61–70. https://doi.org/10.1038/nature12857

18. Passow U, Alldredge AL. Distribution, size, and bacterial colonization of transparent exopolymer particles (TEP) in the ocean. Marine Ecology Progress Series. 1994; 113: 185–198. https://doi.org/10.3354/meps113185

19. Klein C, Claquin P, Pannard A, Napoléon C, Roy BL, Véron B. Dynamics of soluble extracellular polymeric substances and transparent exopolymer particle pools in coastal ecosystems. Marine Ecology Progress Series. 2011; 427: 13–27. https://doi.org/10.3354/meps09049

20. Sun CC, Wang YS, Li QP, Yue WZ, Wang YT, Sun FL, Peng YL. Distribution characteristics of transparent exopolymer particles in the Pearl River estuary, China. Journal of Geophysical Research. 2012; 117: G00N17. https://doi.org/10.1029/2012JG001951.

21. Jennings MK, Passow U, Wozniak AS, Hansell DA. Distribution of transparent exopolymer particles (TEP) across an organic carbon gradient in the western North Atlantic Ocean. Marine Chemistry. 2017; 190: 1–12. https://doi.org/10.1016/j.marchem.2017.01.002

22. Ortega-Retuerta E, Sala MM, Borrull E, Mestre M, Aparicio F, Gallisai R, Antequera C, Marrasé C, Peters F, Simó R, Gasol JM. Horizontal and vertical distributions of transparent exopolymer particles (TEP) in the NW Mediterranean Sea are linked to Chlorophyll a and O2 variability. Frontiers in Microbiology. 2017; 7: 1–12. https://doi.org/10.3389/fmicb.2016.02159.

23. Ning XR, Shi JX, Cai YM, Liu CG. Biological productivity front in the Changjiang Estuary and the Hangzhou Bay and its ecological effects. Acta Oceanologica Sinica. 2004; 26(6): 96–106. (in Chinese with English abstract)

24. Shen ZL. A study of the effects of the three gorge project on the distributions and changes of the nutrients in the Changjiang River estuary. OceanologiaetLimnologia Sinica. 1991; 22(6): 540–546. (in Chinese with English abstract)

25. Tang DL, Di BP, Wei GF, Ni IH, Oh IS, Wang SF. Spatial, seasonal and species variations of harmful algal blooms in the South Yellow Sea and East China Sea. Hydrobiologia. 2006; 568: 245–253. https://doi.org/10.1007/s10750-006-0108-1

26. Zhai WD, Dai MH. On the seasonal variation of air-sea CO2 fluxes in the outer Changjiang (Yangtze River) estuary, East China Sea. Marine Chemistry. 2009; 117: 2–10. https://doi.org/10.1016/j.marchem.2009.02.008

27. Zhai WD, Dai MH, Guo XH. Carbonate system and CO2 degassing fluxes in the inner estuary of Changjiang (Yangtze) River, China. Marine Chemistry. 2007; 107: 342–356. https://doi.org/10.1016/j.marchem.2007.02.011

28. Guo XW, Zhang YS, Zhang FJ, Cao QY. Characteristics and flux of settling particulate matter in neritic waters: the southern Yellow Sea and the East China Sea. Deep-Sea Research II. 2010; 57: 1058–1063. https://doi.org/10.1016Zj.dsr2.2010.02.007

29. Guo SJ, Sun J, Zhao QB, Feng YY, Huang DJ, Liu SM. Sinking rates of phytoplankton in the Changjiang (Yangtze River) estuary: A comparative study between Prorocentrum dentatum and Skeletonema dorhnii bloom. Journal of Marine Systems. 2016; 154: 5–14. https://doi.org/10.1016/jjmarsys.2015.07.003

30. Liu SM, Li RH, Zhang GL, Wang DR, Du JZ, Herbeck LS, Zhang J, Ren JL. The impact of anthropogenic activities on nutrient dynamics in the tropical Wenchanghe and Wenjiaohe Estuary and Lagoon system in East Hainan, China. Marine Chemistry. 2005; 125: 49–68. https://doi.org/10.1016/j.marchem.2011.02.003

31. Welschmeyer NA. Fluorometric analysis of chlorophyll-a in the presence of chlorophyll-b and pheopigments. Limnology and Oceanography. 1994; 39: 1985–1992. https://doi.org/10.4319/lo.1994.39.8.1985

32. Utermöhl H. Zur Vervolkommung der quantitativen Phytoplankton – Methodik. Mitteilung Internationale Vereinigung fuer Theoretische unde Amgewandte Limnologie. 1958; 9: 1–38.

33. Bienfang PK. SETCOL–a technologically simple and reliable method for measuring phytoplankton sinking rates. Canadian Journal of Fisheries and Aquatic Sciences. 1981; 38(10): 1289–1294. https://doi.org/10.1139/f81-173

34. Guo SJ, Feng YY, Wang L, Dai MH, Liu ZL, Bai Y, Sun J. Seasonal variation in the phytoplankton community of a continental-shelf sea: the East China Sea. Marine Ecology Progress Series. 2014; 516: 103–126. https://doi.org/10.3354/meps10952.

35. Mari X, Burd A. Seasonal size spectra of transparent exopolymeric particles (TEP) in a coastal sea and comparison with those predicted using coagulation theory. Marine Ecology Progress Series. 1998; 163: 63–76. https://doi.org/10.3354/meps163063.

36. Alldredge AL, Passow U, Haddock H. The characteristics and transparent exopolymer particle (TEP) content of marine snow formed from thecate dinoflagellates. Journal of Plankton Research. 1998; 20(3): 393–406. https://doi.org/10.1093/plankt/20.3.393

37. Engel A. The role of transparent exopolymer particles (TEP) in the increase in apparent particle stickiness during the decline of a diatom bloom. Journal of Plankton Research. 2000; 22(3): 485–497. https://doi.org/10.1093/plankt/22.3.485

38. Prieto L, Ruiz J, Echevarría F, García CM, Bartual A, Gálvez JA, Corzo A, Macías D. Scales and processes in the aggregation of diatom blooms: high time resolution and wide size range records in a mesocosm study. Deep Sea Research II. 2002; 49: 1233–1253. https://doi.org/10.1016/S0967-0637(02)00024-9

39. Prieto L, Navarro G, Cózar A, Echevarría F, García CM. Distribution of TEP in the euphotic and upper mesopelagic zones of the southern Iberian coasts. Deep Sea Research II. 2006; 53(11-13): 1314–1328. https://doi.org/10.1016/j.dsr2.2006.03.009

40. Claquin P, Probert I, Lefebvre S, Veron B. Effects of temperature on photosynthetic parameters and TEP production in eight species of marine microalgae. Aquatic Microbial Ecology. 2008; 51: 1–11. https://doi.org/10.3354/ame01187.

41. Fukao T, Kimoto K, Kotani Y. Effect of temperature on cell growth and production of transparent exopolymer particles by the diatom Coscinodiscus granii isolated from marine mucilage. Journal of Applied Phycology. 2012; 24: 181–186. https://doi.org/10.1007/s10811-011-9666-3

42. Mari X, Torréton JP, Trinh CBT, Bouvier T, Thuoc CV, Lefebvre JP, Ouillon S. Aggregation dynamics along a salinity gradient in the Bach Dang estuary, North Vietnam. Estuarine, Coastal and Shelf Science. 2012; 96: 151–158. https://doi.org/10.1016/j.ecss.2011.10.028

43. Corzo A, Morillo JA, Rodríguez S. Production of transparent exopolymer particles (TEP) in cultures of Chaetoceros calcitrans under nitrogen limitation. Aquatic Microbial Ecology. 2000; 23: 63–72. https://doi.org/10.3354/ame023063

44. Mari X, Rassoulzadegan F, Brussaard CPD, Wassmann P. Dynamics of transparent exopolymeric particles (TEP) production by Phaeocystis globosa under N- or P-limitation: a controlling factor of the retention/export balance. Harmful Algae. 2005; 4: 895–914. https://doi.org/10.1016/j.hal.2004.12.014

45. Pedrotti ML, Peters F, Beauvais S, Vidal M, Egge J, Jacobsen A, Marrasé C. Effects of nutrients and turbulence on the production of transparent exopolymer particles: a mesocosm study. Marine Ecology Progress Series. 2010; 419: 57–69. https://doi.org/10.3354/meps08840

46. Ramaiah N, Yoshikawa T, Furuya K. Temporal variations in transparent exopolymer particles (TEP) associated with a diatom spring bloom in a subarctic ria in Japan. Marine Ecology Progress Series. 2001; 212: 79–88. https://doi.org/10.3354/meps212079.

47. Corzo A, Rodriguez-Galvez S, Lubian L, Sangrá P, Martínez A, Morillo JA. Spatial distribution of transparent exopolymer particles in the Bransfield Strait, Antarctica. Journal of Plankton Research. 2005; 27(7): 635–646. https://doi.org/10.1093/plankt/fbi038

48. Peng AG, Huang YP. Study on TEP and its relationships with Uranium, Thorium, Polonium Isotopes in Jiulong Estuary. Journal of Xiamen University. 2007; 46(1): 38–42. (in Chinese with English abstract)

49. Wetz MS, Robbins MC, Paerl HW. Transparent exopolymer particles (TEP) in a river-dominated estuary:spatial-temporal distributions and an assessment of controls upon TEP formation. Estuaries and Coasts. 2009;32: 447–455. https://doi.org/10.1007/s12237-009-9143-2

50. Sun CC, Wang YS, Wu ML, Li N, Lin L, Song H, Wang YT, Deng C, Peng YL, Sun FL, Li CL. Distribution of transparent exopolymer particles in the Pearl River Estuary in summer. Journal of Tropical Oceanography. 2010; 29(5): 81–87. (in Chinese with English abstract)

51. Malpezzi MA, Sanford LP, Crump BC. Abundance and distribution of transparent exopolymer particles in the estuarine turbidity maximum of Chesapeake Bay. Marine Ecology Progress Series. 2013; 486: 23–35. https://doi.org/10.3354/meps10362

52. Parinos C, Gogou A, Krasakopoulou E, Lagaria A, Giannakourou A, Karageorgis AP, Psarra S. Transparent exopolymer particles (TEP) in the NE Aegean Sea frontal area: seasonal dynamics under the influence of Black Sea water. Continental Shelf Research. 2017; 149: 112–123. https://doi.org/10.1016/j.csr.2017.03.012

53. Javier R, Carlos MG, Jaime R. Sedimentation loss of phytoplankton cells from the mixed layer: effects of turbulence levels. Journal of Plankton Research. 1996; 18(9): 1727–1734. https://doi.org/10.1093/plankt/18.9.1727

54. Ruiz J, Macías D, Peters F. Turbulence increases the average settling velocity of phytoplankton cells. Proceeding of the National Academy of Science of the United States of America. 2004; 101(51): 17720–17724. https://doi.org/10.1073/pnas.0401539101

55. Azetsu-Scott K, Passow U. Ascending marine particles: significance of transparent exopolymer particles (TEP) in the upper ocean. Limnology and Oceanography. 2004; 49(3): 741–748. https://doi.org/10.4319/lo.2004.49.3.0741

56. Mari X. Does ocean acidification induce an upward flux of marine aggregates? Biogeosciences Discussion. 2008; 5: 1631–1654. https://doi.org/10.5194/bg-5-1023-2008

57. Vicente ID, Ortega-Retuerta E, Romera O, Moreles-Baquero R, Reche I. Contribution of transparent exopolymer particles to carbon sinking flux in an oligotrophic reservoir. Biogeochemistry. 2009; 96: 13–23. https://doi.org/10.1007/s10533-009-9342-8

58. Rochelle-Newall EJ, Mari X, Pringault O. Sticking properties of transparent exopolymeric particles (TEP) during aging and biodegradation. Journal of Plankton Research. 2010; 32(10): 1433–1442. https://doi.org/10.1093/plankt/fbq060

59. Mari X, Passow U, Migon C, Burd AB, Legendre L. Transparent exopolymer particles: effects on carbon cycling in the ocean. Progress in Oceanography. 2017; 151: 13–37. https://doi.org/10.1016/j.pocean.2016.11.002

60. Obernosterer I, Herndl GJ. Phytoplankton extracellular release and bacterial growth: dependence on inorganic N: P ratio. Marine Ecology Progress Series. 1995; 116: 247–257. https://doi.org/10.3354/meps116247

61. Turner JT. Zooplankton fecal pellets, marine snow and sinking phytoplankton blooms. Aquatic Microbial Ecology. 2002; 27: 57–102. https://doi.org/10.3354/ame027057

62. Turner JT. Zooplankton fecal pellets, marine snow, phytodetritus and the ocean’s biological pump. Progress in Oceanography. 2015; 130: 205–248. https://doi.org/10.1016/j.pocean.2014.08.005

